# Mechanistic Insights into 2-5(H)-Furanone-Mediated Inhibition of Angiogenesis Using HUVECs and Zebrafish Models

**DOI:** 10.64898/2026.03.30.715228

**Authors:** Amulya Vijay, Sambhavi Bhagavatheeswaran, Anandan Balakrishnan

## Abstract

Angiogenesis, the process by which new blood vessels form from existing vasculature, is fundamental to tissue repair and regeneration but also underlies pathological conditions such as cancer progression. Targeting angiogenesis has thus become a promising approach for developing novel cancer therapeutics. While various phytochemicals have demonstrated anti-angiogenic effects, the role of 2-5(H)-Furanone, a naturally occurring lactone found in various plants and marine sources with diverse biological activities, remains insufficiently explored. In this study, we systematically evaluate the anti-angiogenic potential of 2-5(H)-Furanone using Human Umbilical Vein Endothelial Cells (HUVECs) as an *in vitro* model and zebrafish embryos as an *in vivo* model.

Experimental findings demonstrated that treatment of HUVECs with increasing concentrations of 2-5(H)-Furanone led to significant, dose-dependent reductions in proliferation, invasion, migration, and tube formation. Analyses of gene expression revealed marked downregulation of key pro-angiogenic mediators, VEGF, and HIF-1α. Complementing these *in vitro* results, *in vivo* studies in zebrafish embryos showed robust, dose-dependent inhibition of intersegmental vessel (ISV) formation, accompanied by suppression of critical angiogenesis-related genes. Molecular docking further supported these observations by indicating stable binding of 2-5(H)-Furanone to major angiogenic targets, including VEGFR2, MMP2, HIF-1α, and PIK3CA.

Collectively, our data demonstrate that 2-5(H)-Furanone potently inhibits angiogenesis, as evidenced in both HUVEC and zebrafish models, through functional and molecular mechanisms. These findings support the further development of 2-5(H)-Furanone as a promising anti-angiogenic therapy candidate.

## INTRODUCTION

Angiogenesis is a precisely coordinated biological process in which new blood vessels form from existing ones through the activation, proliferation, and directed migration of endothelial cells (Dudley & Griffioen, 2023). This process plays a crucial role in tumour metastasis by facilitating the delivery of oxygen and nutrients to growing tumours and enabling tumour cells to enter the circulation, thereby invading distant tissues (Leone et al., 2024). The Folkman hypothesis posits that inhibiting angiogenesis can restrain tumour growth, pushing tumours into a dormant state. Consequently, targeting tumour vasculature through angiogenesis inhibition is considered a promising approach for cancer treatment (Bagley, 2016; Folkman, 1997). This process is vital for various physiological conditions, including embryonic development, wound healing, and tissue repair (Li et al., 2022).

Phytochemicals, which are naturally derived plant compounds, have been recognized for their capacity to modulate angiogenic pathways by affecting multiple molecular signalling mechanisms that govern new blood vessel formation (Janani et al., 2019). 2(5H)-Furanone is a small heterocyclic compound with a five-membered lactone ring, serving as the core structure for many bioactive molecules (Byczek-Wyrostek et al., 2018). It exhibits versatile chemical reactivity and exists in tautomeric forms, enhancing its synthetic utility. Found in natural products, 2(5H)-furanone and its derivatives demonstrate diverse biological activities, including antimicrobial, anticancer, and importantly, anti-angiogenic effects (Alam et al., 2008). Its ability to inhibit angiogenesis, a key process in tumour growth, makes it a promising compound for cancer research and therapeutic development. Some evidence suggests related compounds may influence cell proliferation and migration, but comprehensive studies validating the anti-angiogenic activity of 2-5H-Furanone remain scarce (Park et al., 2008; Zhong et al., 2011).

*In vitro* cell models serve as fundamental tools in cellular and molecular research, aiding target validation and drug development. Human umbilical vein endothelial cells (HUVECs) are widely utilized to investigate angiogenesis under controlled conditions, making them an ideal system for assessing the anti-angiogenic potential of phytochemicals (Zheng et al., 2009).

The zebrafish (Danio rerio) offers an advantageous *in vivo* model for drug discovery and target validation due to its rapid embryonic development and transparent body, which allows direct observation of vascular growth and morphology (Gore et al., 2012). Approximately 24 hours post fertilization (hpf), zebrafish embryonic circulation commences, and the intersegmental vessels (ISVs) emerge from the dorsal aorta and posterior cardinal vein via angiogenic processes (Eberlein et al., 2021). This model is well-suited for angiogenesis studies, owing to its ease of maintenance and permeability to small molecules.

The present study aims to evaluate the anti-angiogenic potential of 2-5H-Furanone employing both HUVEC-based *in vitro* assays and zebrafish embryo *in vivo* models. Furthermore, it seeks to elucidate the molecular mechanisms involved by examining changes in angiogenesis-related gene expression.

## MATERIALS AND METHODS

### Chemicals and Reagents

2-5(H)-Furanone (purity ≥98%), Endothelial growth medium (EGM-2), fetal bovine serum (FBS), penicillin-streptomycin, Matrigel (growth factor-reduced), phosphate-buffered saline (PBS), Ethanol, dimethyl sulphoxide (DMSO), paraformaldehyde (PFA) and sodium acetate were purchased from Sigma Aldrich, USA.

### Cell culture and maintenance

HUVECs (Lonza, Mumbai, India) were cultured in endothelial cell growth medium-2 (EGM-2) BulletKit, consisting of endothelial basal medium (EBM-2) supplemented with essential growth factors and components including ascorbic acid, fetal bovine serum (FBS), insulin-like growth factor (IGF), human epidermal growth factor (hEGF), human fibroblast growth factor (hFGF), and vascular endothelial growth factor (VEGF). Cells at passage number 4 were used for all experiments. Cultures were maintained at 37°C in a humidified incubator with 5% CO2.

### Cell viability assay

Cell viability was evaluated using the Cell Counting Kit-8 (CCK-8; Sigma-Aldrich, USA), a colorimetric assay based on the reduction of WST-8 [2-(2-methoxy-4-nitrophenyl)-3-(4-nitrophenyl)-5-(2,4-disulfophenyl)-2H-tetrazolium, monosodium salt] by cellular dehydrogenases to produce a water-soluble orange formazan dye. HUVECs were seeded at a density of 1 × 10^4^ cells per well in 96-well plates and incubated until reaching 70–80% confluence. Cells were then treated with various concentrations of 2-5(H)-Furanone (2-100 µM) for 24 hours. Following treatment, cell viability was assessed according to the manufacturer’s protocol. Absorbance was measured at 450 nm using a Synergy HT Multi-Mode Microplate Reader (Biotek, USA). Viability was expressed as a percentage relative to untreated control cells.

### Invasion assay

The invasive capability of HUVECs was assessed using Transwell chambers with polycarbonate membrane filters of 8μm pore size (Costar, Corning Incorporated, USA). The inserts were coated with 2mg/ml Matrigel matrix (Cultrex, Trevigen), serving as a reconstituted basement membrane. The lower chamber was filled with 600 μL of endothelial growth medium containing various concentrations of the test compound (e.g., 2-5(H)-Furanone). HUVECs were harvested, resuspended, and seeded into the upper chamber at a density of 1 × 10^4 cells per well in 100 μl medium. The cells were allowed to invade at 37°C for 16 hours. After incubation, the non-invaded cells remaining on the upper surface of the membrane were gently removed with a cotton swab. The inserts were then fixed in 70% ethanol and stained with 0.2% crystal violet. Invaded cells on the lower surface of the membrane were visualized and counted in multiple fields under an inverted microscope, quantifying the degree of invasion in response to the compound.

### Wound-Healing Assay

Wound healing migration assays were conducted using the ibidi CultureInsert 2 Well in a μ-Plate 24 Well format. HUVECs were seeded into each compartment of the Culture-Insert at a density of 35,000 cells per well. Once cells reached approximately 90% confluency, the insert was carefully removed to create a defined cell-free gap (“wound”). Cultures were then incubated with fresh medium containing various concentrations of the test compound 2-5(H)-Furanone. Microscopic images of the wound area were acquired immediately (0 h) and after 24 h incubation at 37°C. The extent of cell migration into the wound area was quantified using ImageJ software by measuring wound closure, allowing for assessment of the compound’s effect on endothelial cell motility.

### Tube Formation Assay

The tube-forming ability of HUVECs was evaluated using the Cultrex® *In vitro* Angiogenesis Assay Tube Formation Kit (Trevigen, Gaithersburg, MD, USA). Briefly, 50μl of basement membrane extract (BME) solution was added to each well of a 96-well plate and incubated at 37°C for 1 hour to allow gel formation. HUVECs were seeded onto the solidified BME at a density of 3 × 10^3^ cells per well in the presence of various concentrations of the test compound. All treatments were performed in medium containing 20ng/ml VEGF. Sulforaphane at 5 μM served as a positive control, while 20 ng/ml VEGF alone served as a negative control. Plates were incubated for 4–6 hours at 37°C until capillary-like tube networks formed. After incubation, the media was aspirated, and 100 μL of 2 mM calcein AM was added for 30 minutes to stain viable cells. Tube networks were visualized using a fluorescence microscope at 10× magnification, and images were obtained from multiple fields. Quantitative analysis of tube networks (e.g., total tube length and branch points) was performed using ImageJ with the Angiogenesis Analyzer plugin.

### Quantitative Real-Time PCR (qRT-PCR)

Total RNA was extracted from HUVECs treated with various concentrations of isopimpinellin using RNAzol® RT reagent (Sigma Aldrich, USA) following the manufacturer’s instructions. RNA concentration and purity were measured using a NanoDrop 2000 spectrophotometer (Thermo Scientific, Germany). For mRNA analysis, 500 ng of total RNA was reverse-transcribed into complementary DNA (cDNA) using the Thermo Revert-aid First Strand cDNA Synthesis Kit (Thermo Scientific, USA). Quantitative PCR amplification for mRNA was performed using GoTaq® qPCR Master Mix (Promega). Relative gene expression levels were calculated using the 2^−ΔΔCt method. Primer sequences used in this study are listed in Supplementary Tables 1.

### *In vivo* Experiments

#### Zebrafish Maintenance and Breeding

Adult wild-type zebrafish (Danio rerio) were procured from a local supplier and maintained under standardized conditions to preserve genetic diversity. The fish were housed in glass tanks with continuous aeration at 28°C. Water quality was maintained by replacing the water with fresh reverse osmosis water every two days. The fish were subjected to a controlled photoperiod of 14 hours of light and 10 hours of darkness. Natural pairwise mating was used to produce embryos, which were raised in embryo medium under standard laboratory conditions.

#### Drug Treatment and Angiogenesis Assessment

At 24 hours post-fertilization (hpf), zebrafish embryos were carefully dechorionated and exposed to various concentrations of 2-5(H)-Furanone (0–200 μM) dissolved in E3 medium. Based on preliminary viability tests, appropriate doses were selected for further analysis. Treated embryos were incubated with 2-5(H)-Furanone for 48 hours starting at 24 hpf. For wild-type embryos, after treatment, embryos were fixed with 4% paraformaldehyde (PFA), stained with o-dianisidine solution, and cleared for visualization of intersegmental vessels (ISVs). For the transgenic line Tg(hspGFFDMC84A), treatment consisted solely of drug exposure followed by incubation, after which larvae were directly photographed to assess vascular development using a Leica M165FC fluorescence stereo microscope. Representative images were captured for documentation and analysis.

#### Quantitative Real-Time PCR (qRT-PCR)

Total RNA was extracted from pools of 50 zebrafish embryos per treatment group. Embryos were homogenized using a micropestle in RNAzol® RT reagent (Sigma-Aldrich, USA) according to the manufacturer’s protocol. RNA purity and concentration were measured using a NanoDrop 2000 spectrophotometer (Thermo Scientific, Germany). For cDNA synthesis, 500 ng of total RNA was reverse transcribed using the Thermo Revert-aid First Strand cDNA Synthesis Kit (Thermo Scientific, USA). Quantitative PCR was performed using GoTaq® qPCR Master Mix (Promega) to amplify target angiogenesis-related genes. Data were analyzed using the comparative 2^(-ΔΔCt) method to determine relative gene expression. The primer sequences used in the study are listed in Supplementary Table 2.

#### Statistical Analysis

Data analysis and graphical presentations were performed using GraphPad Prism version 5.0 (GraphPad Software, USA). All experimental procedures were conducted in triplicate. Results are presented as mean values ± standard error of the mean (SEM). Statistical significance among groups was evaluated using one-way analysis of variance (ANOVA) followed by the Newman-Keuls post hoc test. Significance levels are denoted as ***P < 0.001, **P < 0.01, and *P < 0.05.

## RESULTS

### Chemical Structure and Cytotoxicity of 2-5(H)-Furanone

The chemical structure of 2-5(H)-Furanone, a five-membered oxygen-containing lactone ring compound, is depicted in Figure 1. Cytotoxic evaluation using HUVECs revealed that 2-5(H)-Furanone significantly inhibited cell viability in a dose-dependent manner after 24 hours of treatment. The calculated half-maximal inhibitory concentration (IC50) was 16.34 μM, confirming the compound’s potent cytotoxic effect on endothelial cells. This cytotoxicity suggests that 2-5(H)-Furanone can effectively inhibit endothelial proliferation, an essential process in angiogenesis.

**Figure 1.**
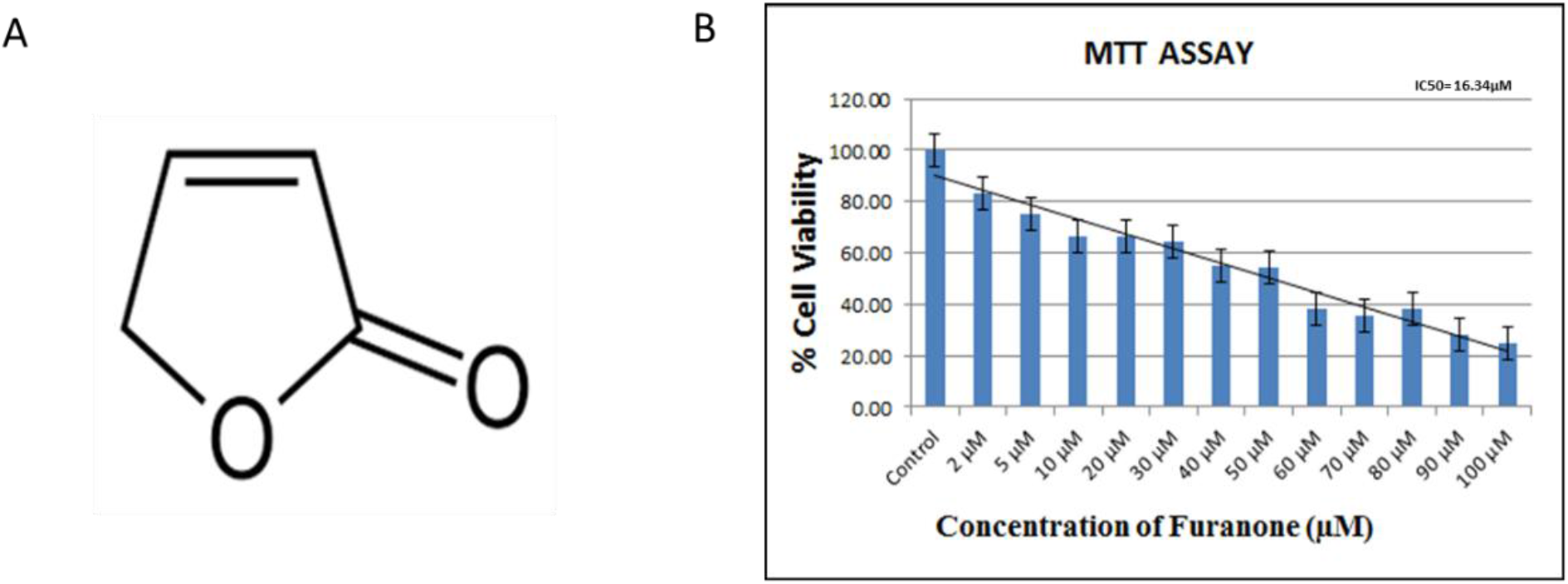
**(A)** Chemical structure of 2-5(H)-Furanone **(B)** bar graph presenting HUVEC viability (%) following 24-hour treatment with varying concentrations of 2-5(H)-Furanone, measured by CCK-8 assay. Data are mean ± SEM. The graph indicates a dose-dependent decline in viability.

### 2-(5)H-Furanone Inhibits HUVEC Invasion

Exposure of HUVECs to incremental concentrations of 2-5(H)-Furanone significantly reduced their invasive capacity through Matrigel-coated transwell membranes in a clear dose-dependent manner (Figure 2). This assay models endothelial cell remodelling and migration through extracellular matrix components, a prerequisite for angiogenic sprouting and new vessel formation. Reduced invasion indicates that 2-5(H)-Furanone may interfere with endothelial extracellular matrix degradation or motility, potentially impairing angiogenesis.

**Figure 2.**
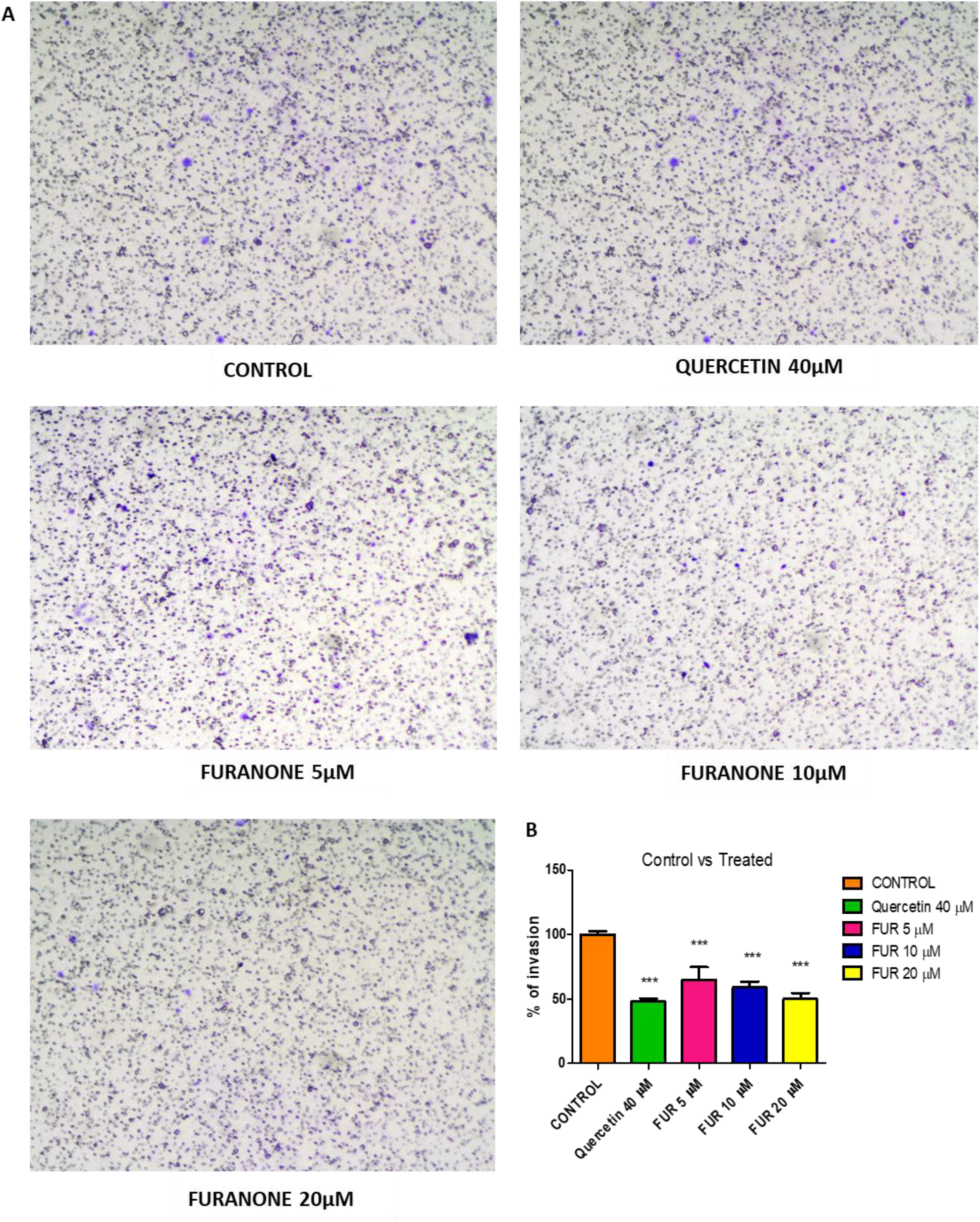
**(A)** Representative images of HUVECs that have invaded through the transwell membrane after exposure to different concentrations of 2-5(H)-Furanone (magnification ×10), **(B)** corresponding quantification graph displaying the percentage of invasive cells relative to control. Data are mean ± SEM; ***P < 0.001 compared to control.

### 2-5(H)-Furanone Suppresses Endothelial Cell Migration

The wound-healing assay demonstrated that 2-5(H)-Furanone progressively inhibited HUVEC migration into the scratched gap over 24 hours (Figure 3A). Quantitative image analysis showed a significantly higher percentage of unclosed wound area in treated cells compared to vehicle controls, reinforcing the inhibitory effect on cell motility. This result suggests that 2-5(H)-Furanone disrupts cellular processes critical for endothelial migration, a key event in vascular remodelling and repair.

**Figure 3.**
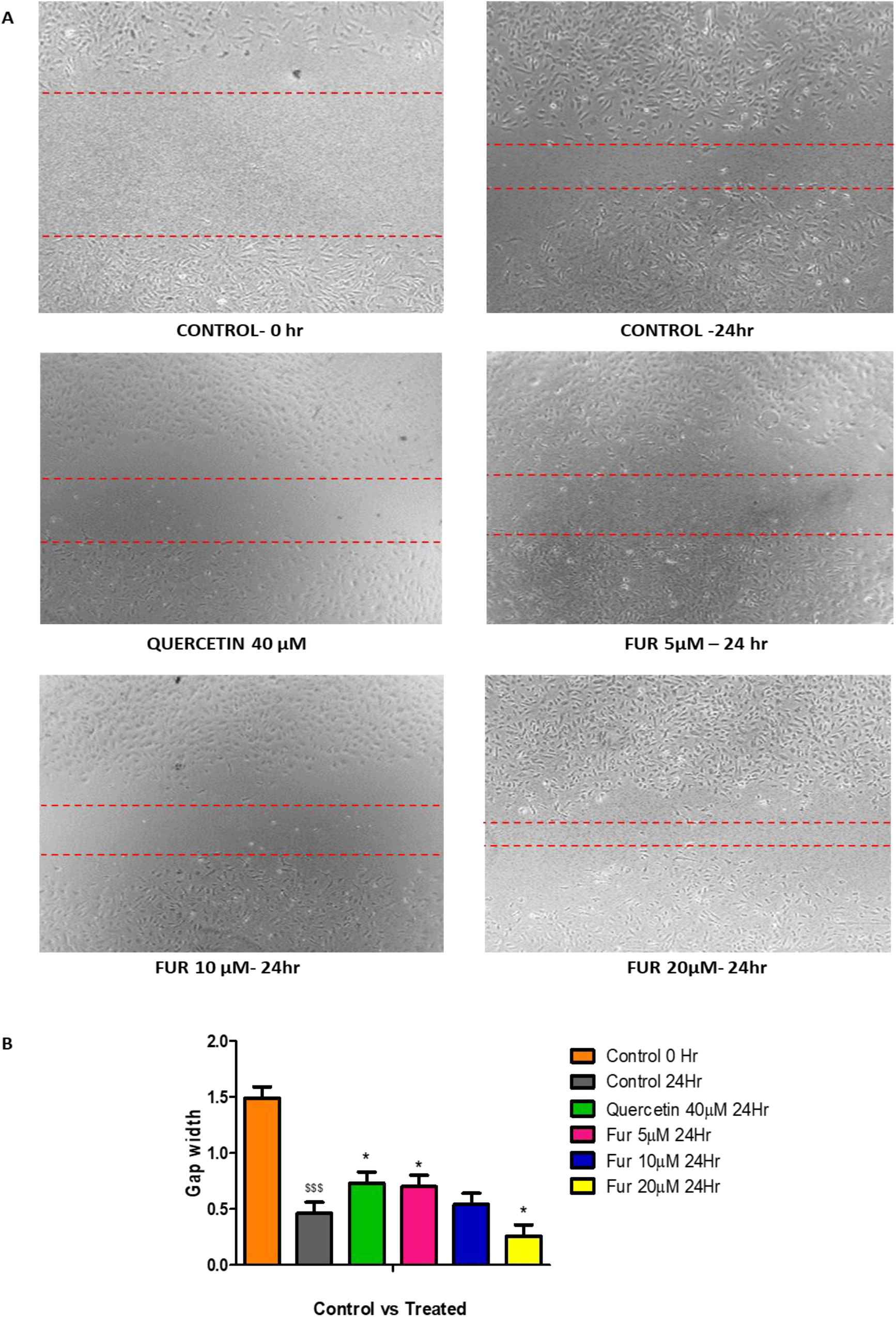
**(A)** Representative phase-contrast images of wound area closure at 0- and 24-hours post-scratch after treatment with vehicle, quercetin (positive control), or various concentrations of 2-5(H)-Furanone (magnification ×4), **(B)** graph shows the percentage of the gap width remaining unclosed after 24 hours. Data represent mean ± SEM; ***P < 0.001, *P < 0.05 versus control

### Disruption of Tube Formation Ability by 2-5(H)-Furanone

Treatment with 2-5(H)-Furanone markedly impaired HUVECs’ ability to form capillary-like tubular structures on Matrigel. Objective quantification revealed significant reductions in total tube length, number of branch points, and loop formation compared to controls (Figure 4A). Since tube formation *in vitro* correlates closely with angiogenic capacity *in vivo*, this finding underlines the compound’s potent anti-angiogenic activity at the cellular morphogenesis stage.

**Figure 4.**
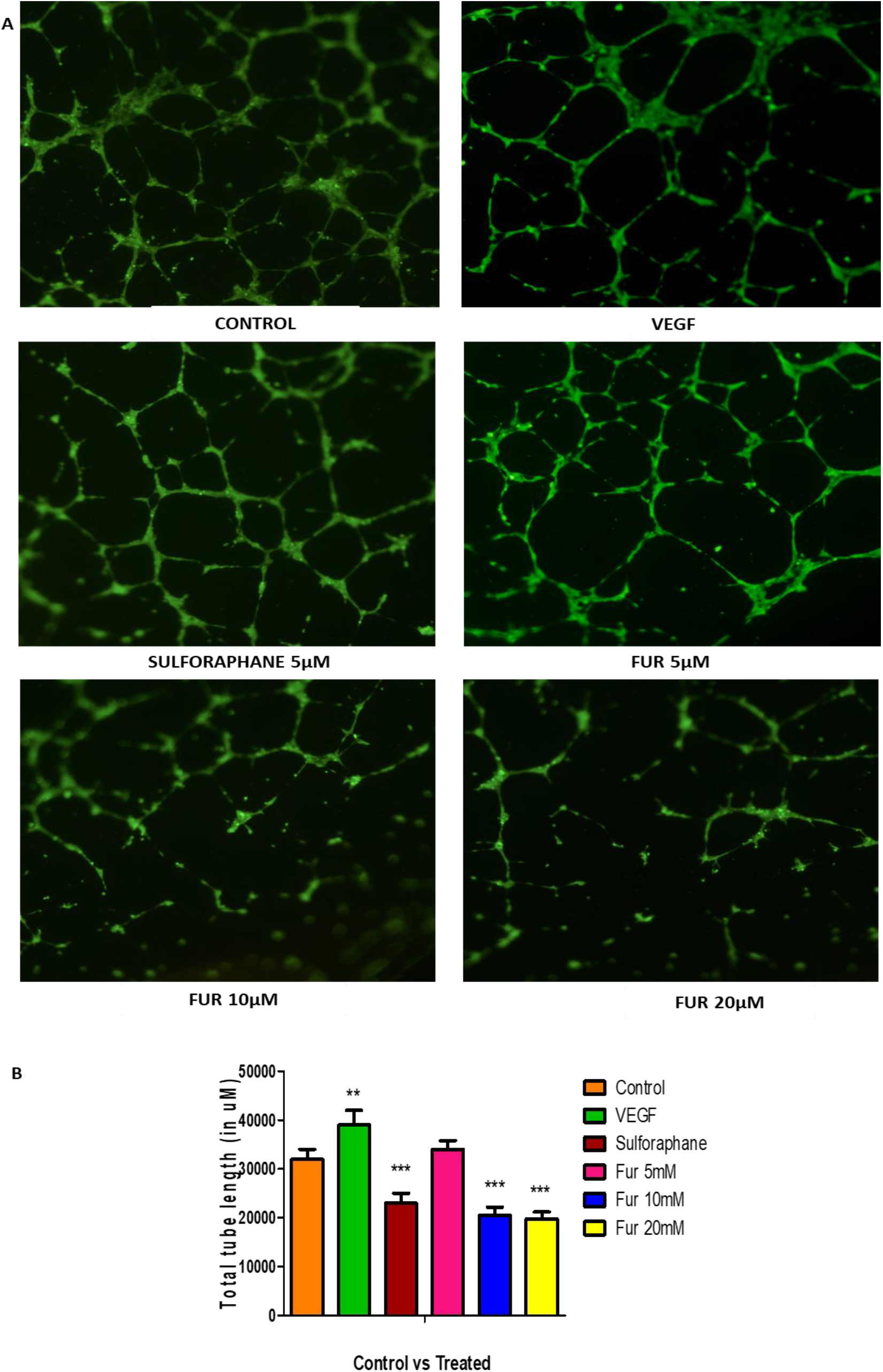
**(A)** Representative phase-contrast microscopy images of tubular network formation by HUVECs treated with vehicle or 2-5(H)-Furanone (magnification ×4). **(B)** Quantitative analysis of total tube length, measured in micrometers, showing significant dose-dependent reductions. Data are mean ± SEM; ***P < 0.001 compared to control.

### Downregulation of Pro-Angiogenic Gene Expression by 2-5(H)-Furanone

qRT-PCR analysis revealed dose-dependent decreases in the mRNA expression of pro-angiogenic mediators *VEGF* and *HIF-1α* in HUVECs treated with 2-5(H)-Furanone (Figure 5). *VEGF* is a critical growth factor promoting endothelial proliferation and migration, whereas *HIF-1α* regulates hypoxia-induced *VEGF* expression. These decreases suggest molecular inhibition of canonical angiogenic signalling pathways that contribute to vessel formation and survival.

**Figure 5.**
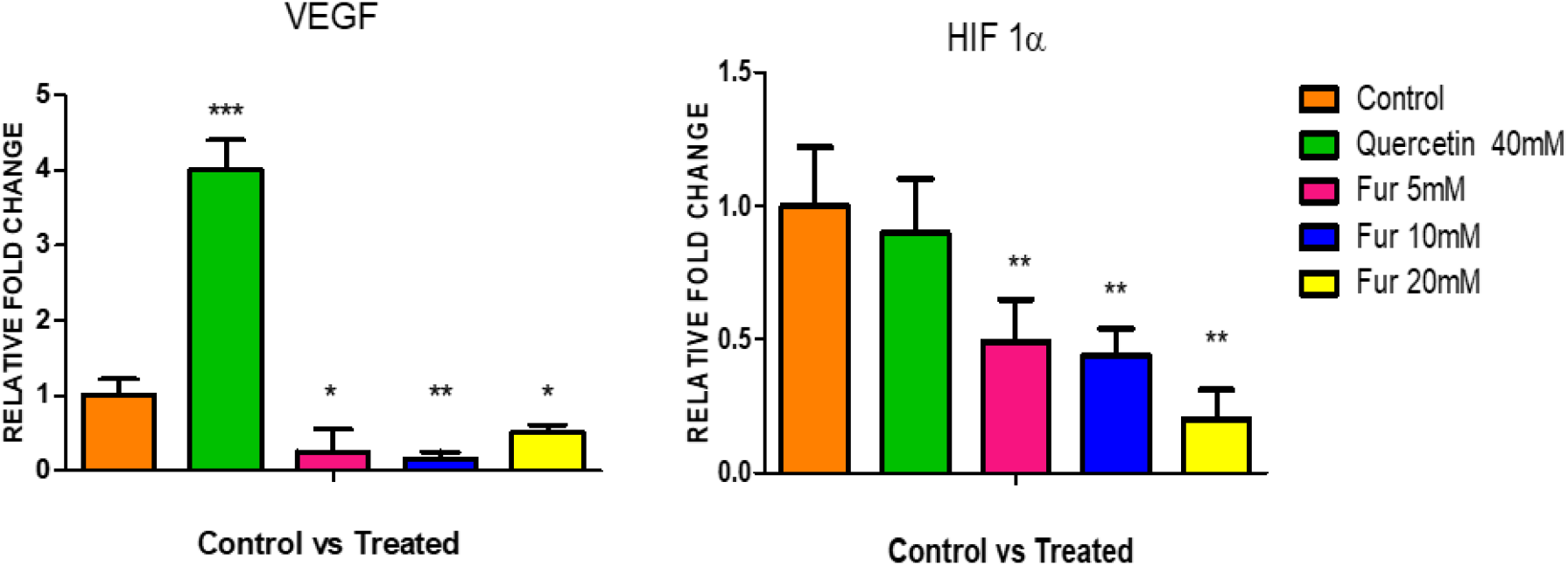
Relative mRNA expression levels of VEGF and HIF-1α determined by qRT-PCR in HUVECs after treatment with increasing concentrations of 2-5(H)-Furanone. Expression levels are normalized to GAPDH. Data shown as mean ± SEM; ***P < 0.001, **P < 0.01, *P < 0.05 versus control.

**Figure 6.**
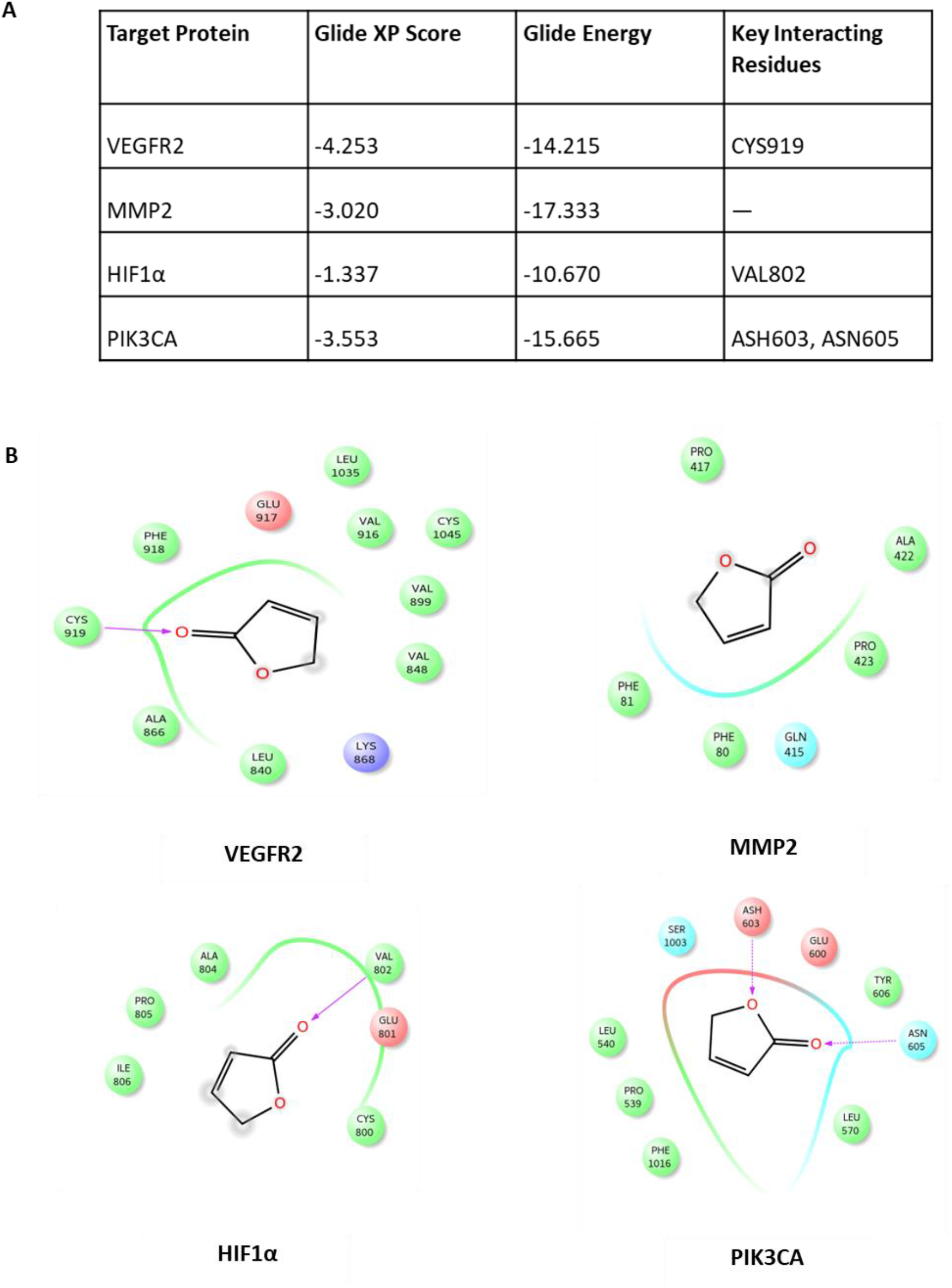
**(A)** Docking results table showing Glide XP scores, Glide energies, and **(B)** key interacting residues of 2-5(H)-Furanone with angiogenesis-related targets VEGFR2, MMP2, HIF1α, and PIK3CA.

### Molecular Docking of 2-5(H)-Furanone with Angiogenesis-Related Targets

In silico docking studies demonstrated that 2-5(H)-Furanone stably associates with key angiogenesis-related proteins, including VEGFR2, MMP2, HIF-1α, and PIK3CA. The favourable glide scores and key ligand-protein interactions suggest that the compound may directly inhibit these proteins’ function, disrupting pathways essential for angiogenesis such as receptor tyrosine kinase signalling, matrix remodelling, hypoxia response, and PI3K/AKT-mediated survival.

### 2-5(H)-Furanone Inhibits Intersegmental Vessel (ISV) Formation in Zebrafish Embryos

*In vivo* assessment using the zebrafish embryo model revealed that treatment with 2-5(H)-Furanone caused progressive dose-dependent inhibition of ISV development, as visualized by o-dianisidine staining of red blood cells (Figure 7). Additional experiments using the transgenic zebrafish line Tg(hspGFFDMC84A) demonstrated that drug treatment similarly disrupted vascular development (Figure 8), as confirmed by fluorescence imaging under a Leica M165FC stereo microscope. These findings validate the anti-angiogenic potential of 2-5(H)-Furanone in a whole-organism context, demonstrating its ability to impair vascular formation under physiologically relevant conditions.

**Figure 7.**
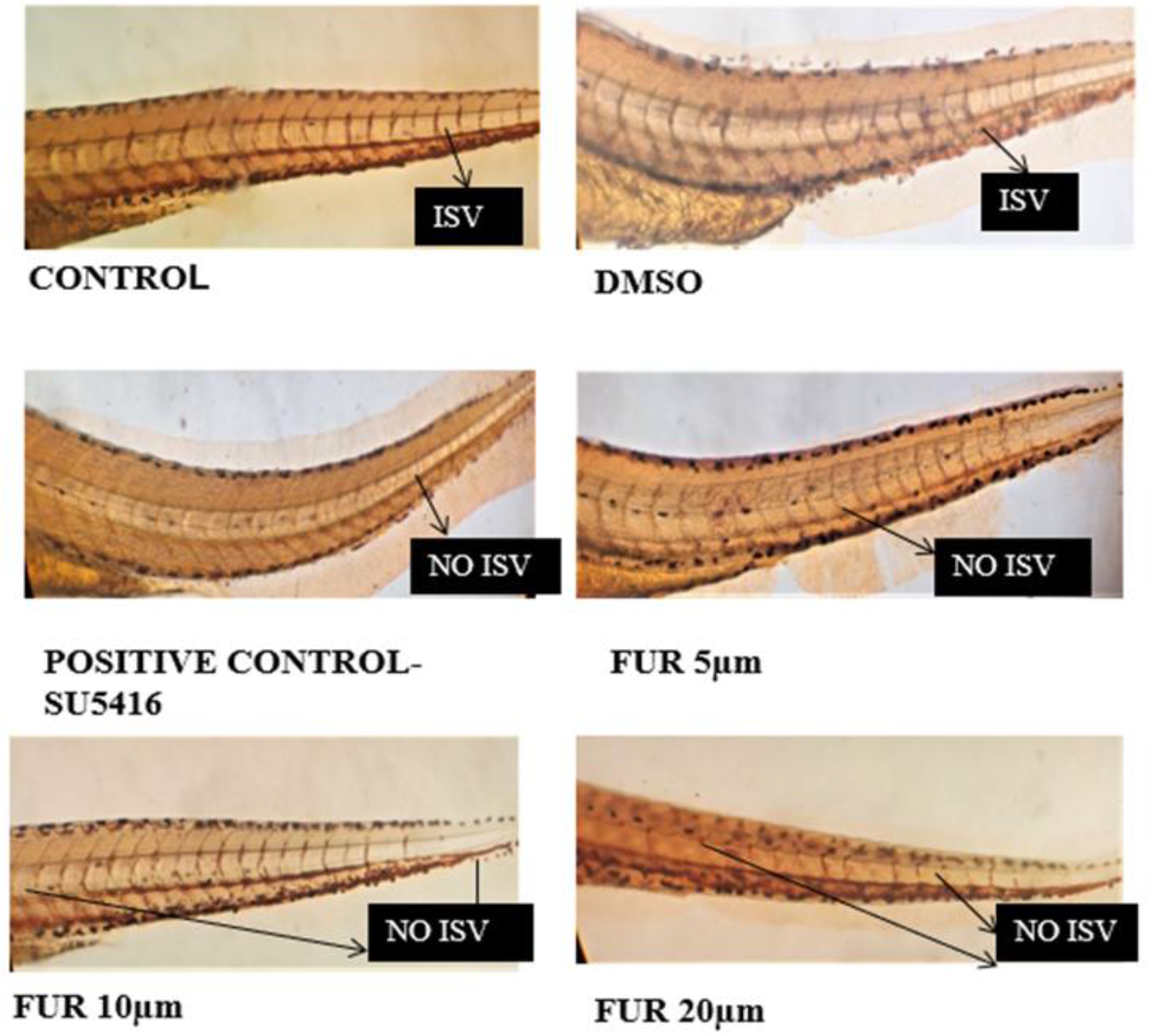
RBC staining images of control, 0.3% DMSO, positive control, and zebrafish embryos treated with various concentrations of 2-5(H)-Furanone, viewed at 10× magnification. Reductions in ISV formation are indicated by decreased staining and disrupted vessel patterning

**Figure 8.**
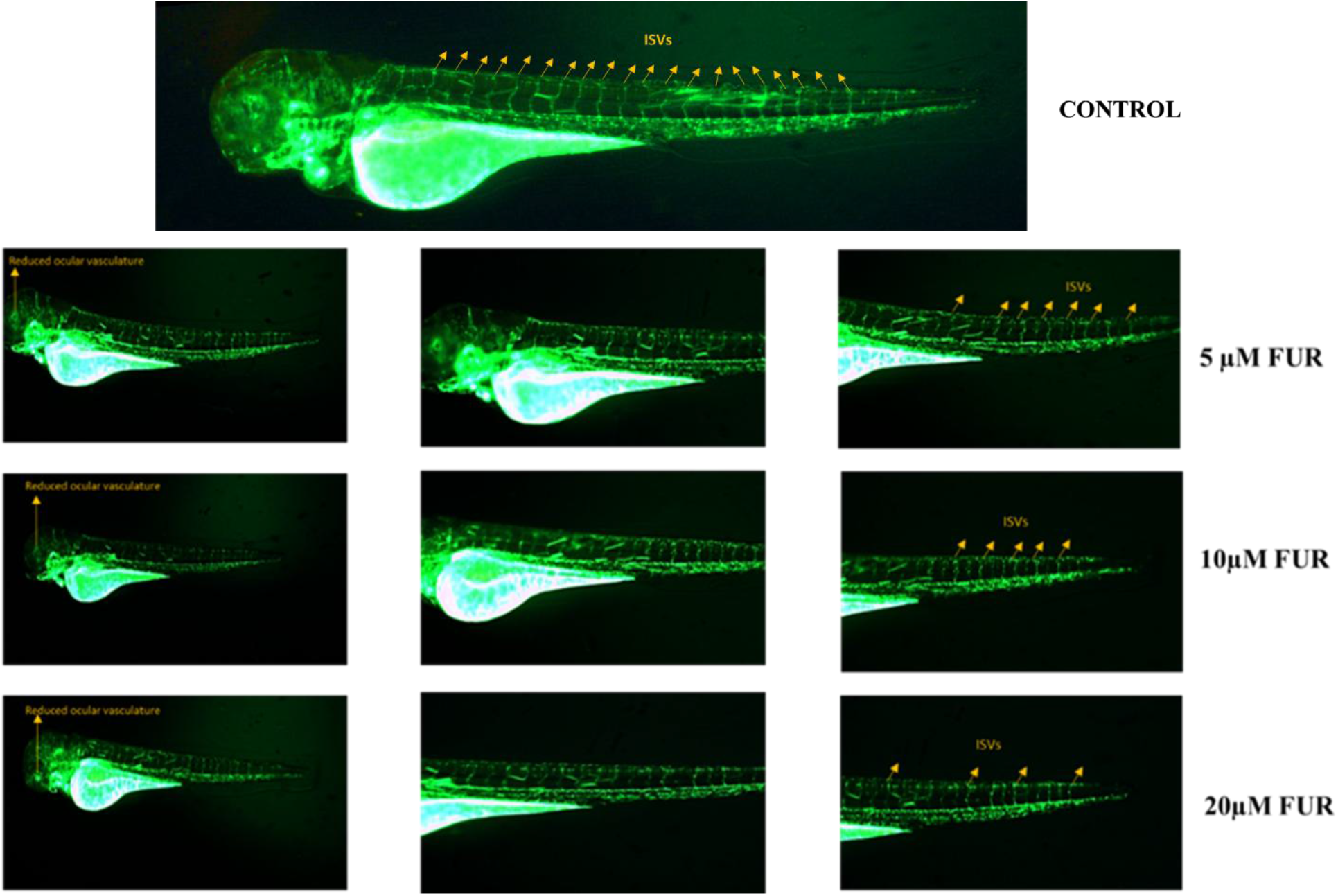
Dose-dependent response of ISV formation in transgenic Tg(hspGFFDMC84A) zebrafish embryos treated with 2-5(H)-Furanone, showing progressive inhibition with increasing drug concentrations.

### Downregulation of Angiogenesis-Related Genes in Zebrafish Embryos

qRT-PCR profiling of treated zebrafish embryos showed significant dose-dependent suppression of key angiogenic genes (Figure 9), including *VEGF, VEGFR2, SURVIVIN, ANGPT1, ANGPT2, TIE1*, and *TIE2*. These genes regulate vessel growth, maturation, and survival signaling, and their downregulation correlates with the impaired ISV formation observed. This cross-species molecular effect supports the conserved anti-angiogenic mechanisms of 2-5(H)-Furanone.

**Figure 9.**
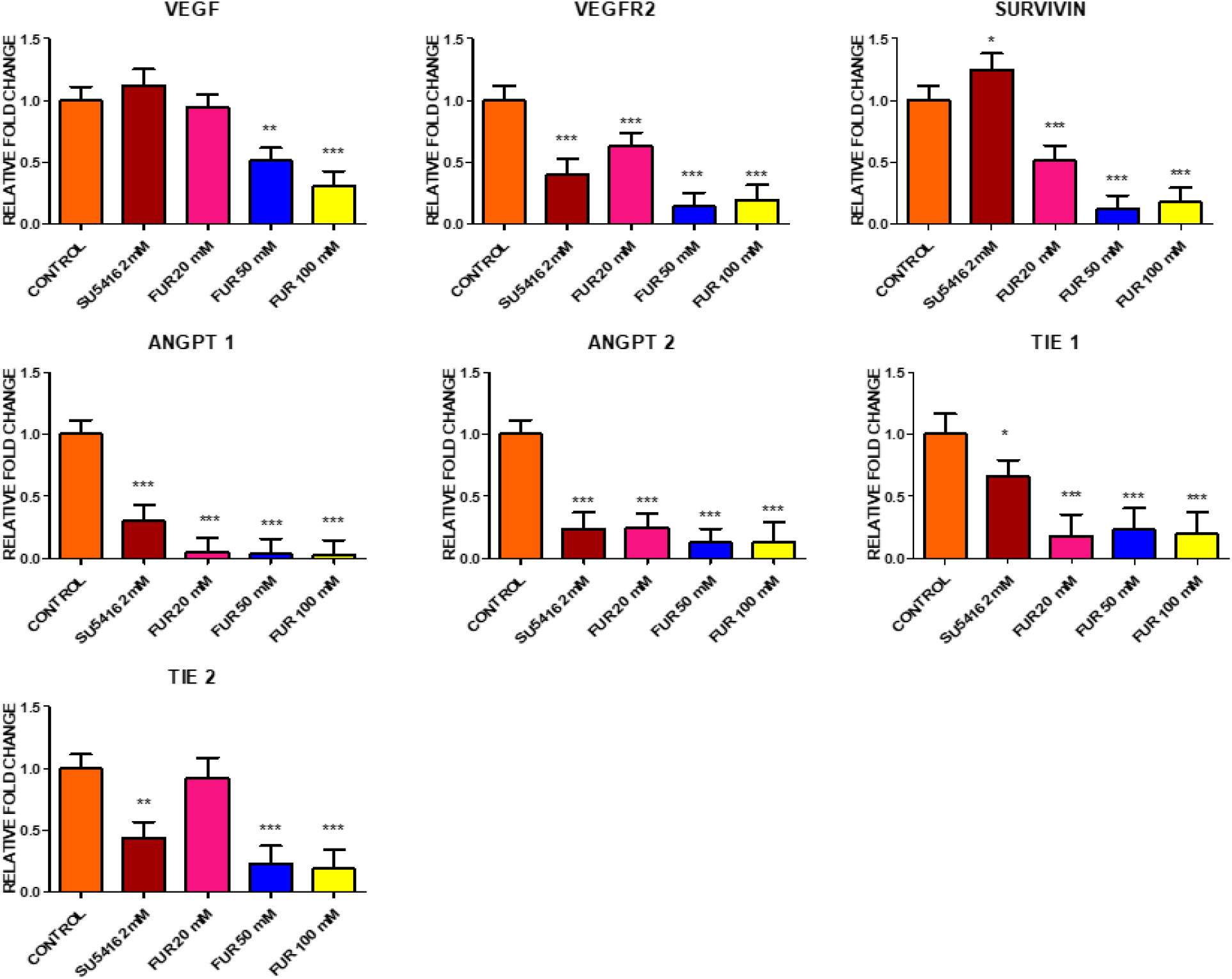
Gene expression profiling of angiogenic markers in control and treated zebrafish embryos after 2-5(H)-Furanone exposure. Relative fold changes were determined by qRT-PCR. Statistical significance denoted as ***P < 0.001, **P < 0.01, and *P < 0.05 compared to control.

## DISCUSSION

Angiogenesis is a tightly regulated biological process essential for normal physiological processes, such as tissue repair, and for pathological conditions, including tumour development and metastasis. Consequently, targeting angiogenesis has emerged as a promising therapeutic strategy for cancer and other angiogenesis-dependent diseases (Akbarian et al., 2022; Liu et al., 2025; Teleanu et al., 2019). This study explored the anti-angiogenic potential of 2-5(H)-Furanone, a phytochemical with largely uncharacterized vascular effects. Our findings demonstrate that 2-5(H)-Furanone effectively inhibits several critical steps of angiogenesis, including endothelial cell viability, invasion, migration, and tubulogenesis, accompanied by downregulation of major pro-angiogenic genes, both *in vitro* and *in vivo*.

The dose-dependent reduction in HUVEC viability, with an IC50 of 16.34 μM after 24 hours, suggests that 2-5(H)-Furanone exerts significant cytotoxicity against endothelial cells. This cytotoxicity is particularly relevant given the importance of endothelial proliferation in supporting neovascularization during tumour progression. Similar anti-endothelial effects have been reported for other furanone derivatives, which mediate cytostatic and cytotoxic responses by disrupting cell cycle progression and inducing apoptosis (He et al., 2017). Additionally, compounds such as curcumin and epigallocatechin gallate, known for their anti-angiogenic properties, exhibit pronounced cytotoxicity against endothelial cells, positioning 2-5(H)-Furanone within this effective class of phytochemicals (Park et al., 2008).

In terms of invasive capacity, 2-5(H)-Furanone substantially impaired endothelial cells’ ability to penetrate Matrigel-coated membranes, a critical step mimicking extracellular matrix degradation during vessel sprouting. Given that uncontrolled invasion contributes to pathological angiogenesis and tumour metastasis, this inhibition is highly significant. Endothelial invasion facilitates new vessel formation through matrix remodelling, and disrupting this process can effectively hinder angiogenesis (Park et al., 2008; Staton et al., 2009). Our results here are consistent with other phytochemicals, such as resveratrol and thymoquinone, which exhibit anti-angiogenic effects by targeting endothelial migration and matrix-degrading pathways (Amin et al., 2012; Paramasivam et al., 2012; Farooqi et al., 2022).

Alongside invasion, endothelial cell motility is essential for angiogenic sprouting and tissue repair. Our migration assays demonstrated a marked, dose-dependent suppression of HUVEC migration by 2-5(H)-Furanone, as evidenced by reduced wound closure in scratch assays. This finding aligns with observations from other natural compounds, such as quercetin, which inhibit endothelial movement by interfering with cytoskeletal dynamics and cell motility signaling pathways (Ravishankar et al., 2015; Varinská et al., 2018). Impairing migration is a hallmark of anti-angiogenic agents, as they limit cells’ ability to extend vascular networks and repair tissues (García‐Conesa et al., 2008; Tonutti et al., 2010).

Furthermore, the compound significantly disrupted the morphogenic stage of angiogenesis by impairing the assembly of capillary-like tubular structures on Matrigel. Both total tube length and branch points were significantly decreased, underscoring the inhibitory effect on endothelial network formation. This stage typically reflects angiogenic capacity *in vitro* and serves as a key indicator for functional angiogenesis inhibition. Previous studies have demonstrated that furanocoumarins and related natural products interfere with tube formation by suppressing VEGF and modulating extracellular matrix interactions (Kadioglu et al., 2013; Mashreghi et al., 2018). Our data provides the first direct evidence that 2-5(H)-Furanone shares this effect, reaffirming its potential as a multipronged anti-angiogenic compound (Goodwin, 2007).

At the molecular level, quantitative PCR analysis revealed clear downregulation of essential pro-angiogenic genes, including VEGF and HIF-1α, following treatment with 2-5(H)-Furanone. VEGF drives endothelial proliferation, migration, and survival, while HIF-1α acts as a primary hypoxia-responsive transcription factor promoting VEGF expression (Chen et al., 2015; Hasebe et al., 2003). The repression of these genes mechanistically correlates with the functional inhibition observed in cell assays. The secondary angiogenesis-related marker AKT was similarly diminished, implying broad suppression across canonical signaling pathways. These molecular effects are consistent with those of other anti-angiogenic phytochemicals, such as thymoquinone and furanodiene, which also downregulate these critical genes (Paramasivam et al., 2016; Zhong et al., 2017).

Molecular docking studies further illuminated potential direct interactions between 2-5(H)-Furanone and pivotal angiogenesis-related proteins, namely VEGFR2, MMP2, HIF-1α, and PIK3CA. The favourable binding affinities and stable interactions within active sites suggest that 2-5(H)-Furanone may inhibit angiogenesis by interfering with key receptor tyrosine kinases and downstream signaling molecules. This mechanism is supported by pharmacological evidence, where selective inhibition of VEGFR2 and downstream effectors, such as PI3K/AKT, effectively abrogates angiogenesis and tumour growth (Brown et al., 2010; Kadioglu et al., 2013).

Extending these findings to an *in vivo* context, treatment of zebrafish embryos with 2-5(H)-Furanone substantially impaired intersegmental vessel development, confirmed via o-dianisidine staining. The zebrafish model provides a dynamic platform for evaluating angiogenesis within an intact organism, and our results endorse the translational relevance of 2-5(H)-Furanone’s anti-angiogenic action beyond isolated cell systems. Comparable anti-angiogenic effects have been documented with other natural compounds in zebrafish, underscoring the model’s predictive value (Chen et al., 2018; Bhagavatheeswaran et al., 2021; Langheinrich, 2003).

Complementary qRT-PCR analyses in zebrafish embryos revealed dose-dependent downregulation of a suite of angiogenesis-associated genes, including vegf, vegfr2, survivin, angiopoietins (angpt1, angpt2), and Tie receptors (tie1, tie2). The broad-spectrum gene suppression supports impaired vascular development and indicates that 2-5(H)-Furanone targets conserved signalling nodes crucial for vessel formation and maturation, paralleling patterns seen with established anti-angiogenic drugs (Ma et al., 2007; Thurston & Daly, 2012).

Taken together, the results of this study indicate that 2-5(H)-Furanone can modulate angiogenesis by coordinating effects on endothelial cell function and gene expression. While these findings suggest its role as a modulator of angiogenic processes, several limitations should be considered. The mechanistic insights are primarily based on gene expression and in silico analyses and therefore require validation at the protein and signalling pathway levels. Additionally, although the zebrafish model provides valuable *in vivo* insight, further evaluation in mammalian systems will be necessary to assess translational relevance, pharmacokinetics, and safety. Further investigation into its molecular targets and biological effects will be important to better define its role in angiogenesis-related contexts.

## CONCLUSION

2(5H)-Furanone demonstrates significant anti-angiogenic activity by inhibiting key endothelial cell functions essential for neovascularization, including proliferation, migration, and tube formation, while also downregulating the expression of critical pro-angiogenic genes. Mechanistically, molecular docking suggests that its effects may be primarily mediated by interaction with VEGFR2, with potential modulation of the PIK3CA signalling pathway.

Furthermore, its consistent efficacy in both human endothelial cell systems and *in vivo* zebrafish models underscores its potential as a multi-targeted therapeutic candidate for angiogenesis-related disorders, including cancer. However, further preclinical investigations are necessary to validate its safety, specificity, and translational applicability.

## Acknowledgements

This study was funded and supported by the UGC-UPE phase II thrust area “Herbal Sciences and Drug Development – Theme B. The authors also thank DST-FIST, UGC-SAP and DHR-MRU for the infrastructural facility. We sincerely thank Professor Koichi Kawakami, Laboratory of Molecular and Developmental Biology, Department of Biological Sciences, National Institute of Genetics (NIG), Japan, for providing the transgenic zebrafish line and valuable technical support. This work was conducted under the NIG Joint Research Program grant [NIG JOINT (A), Reference No.: 49A2019].

## Author contributions

AV: conceptualization, methodology, investigation, writing-original draft. AB: conceptualization, supervision, writing-review, and editing.

## Ethical approval

Ethical clearance for the study was granted by the Institutional Animal Ethics Committee, with approval number [Approval No.: 01/36/2015].

