## Supplementary data 1 for "Mechanistic Insights into 2-5(H)-Furanone-Mediated Inhibition of Angiogenesis Using HUVECs and Zebrafish Models"

### Supplementary Table 1: Primers used in qRT-PCR of HUVEC mRNA

| Gene | Forward (5' – 3') | Reverse (5' – 3') |
|---|---|---|
| VEGF | CGGCGAAGAGAAGAGACAC | GGAGGAAGGTCAACCACTCA |
| HIF-1α | AACATAAAGTCTGCAACATGGAAG | TTTGATGGGTGAGGAATGGG |
| ACTB | ACCTTCTACAATGAGCTGCG | CCTGGATAGCAACGTACATGG |

### Supplementary Table 2: Primers used in qRT-PCR of Zebrafish mRNA

| Gene | Forward (5' – 3') | Reverse (5' – 3') |
|---|---|---|
| vegf | AAAAGAGTGCGTGCAAGACC | GACGTTTCGTGTCTCTGTCG |
| vegfr2 | GATGGAGATACACACCTTCAG | TGCGTACCGATGACACATTTC |
| survivin | CACTCCAGAAAACATGGCTAAA | CCATCCTTCCAGCTCTTTCA |
| angtpt 1 | CGTCGCGGTTGGAAATTCAG | AGGTCAGATTTCTCCGTCCG |
| angpt 2 | AGGTGGAGGCTGGACTGTC | GTGGTGAGCAGGTGGATGAC |
| tie 1 | GGCTCTTCTGGCCCTCTTTT | TTGGTCGTCGGGTAAGTGTG |
| tie 2 | CTACCCAGTGACCAACGC | GCTCTACAGCTCCTGACGAT |
| β-actin | CGAGCAGGAGATGGGAACC | CAACGGAAACGCTCATTGC |
